# Serotonergic mediation of orienting and defensive responses in zebrafish

**DOI:** 10.1101/2023.11.24.568601

**Authors:** Bianca Gomes do Nascimento, Jeane Rodrigues Rodrigues, Maria Clara Rosa Silva, Bruna Patrícia Dutra Costa, Marissol Leite, Maryana Pereira Pyterson, Hadda Tercya Lima Silva, Loanne Valeria Xavier Bruce de Souza, Diógenes Henrique de Siqueira Silva, Caio Maximino

**Affiliations:** Laboratório de Neurociências e Comportamento “Frederico Guilherme Graeff”, Faculdade de Psicologia, Instituto de Estudos em Saúde e Biológicas, Universidade Federal do Sul e Sudeste do Pará, Marabá/PA, Brazil; Rede de Biotecnologia e Biodiversidade da Amazônia Legal, Marabá/PA, Brazil; Programa de Pós-Graduação em Reprodução Animal na Amazônia (ReproAmazon), Universidade Federal Rural da Amazônia / Universidade Federal do Pará, Belem/PA, Brazil; Grupo de Estudos da Reprodução de Peixes Amazônicos, Faculdade de Ciências Biológicas, Instituto de Estudos em Saúde e Biológicas, Universidade Federal do Sul e Sudeste do Pará, Marabá/PA, Brazil; Programa de Pós-Graduação em Neurociências e Comportamento, Universidade Federal do Pará, Belem/PA, Brazil

**Keywords:** serotonergic tone, visual attention, general arousal, emotion-cognition interface

## Abstract

Serotonin (5-HT) is involved in arousal and defensive responses, both of which represent modulators of attention and orienting. Orienting responses (ORs), considered as a unit of attentional processing, are elicited by novel innocuous stimuli, differ from defense reflexes (DR)s, as the latter are elicited by painful and threatening stimuli. When zebrafish (Danio rerio) are exposed to a conspecific alarm substance (CAS), visual stimuli elicit a DR instead of ORs, suggesting a state of hypervigilance. CAS elicited an OR-to-DR shift, as shown both by the shift from an approach to an escape response when the stimulus was turned on and by the changes in directional focus. Both pCPA and 5-HTP attenuated the CAS-elicited OR-to-DR shift . While pCPA did not alter ORs in animals which were not exposed to CAS, 5-HTP abolished those responses, suggesting that the serotonergic tone is important in regulating the general arousal levels in zebrafish.

## Introduction

Orienting responses (ORs), originally described by Ivan Pavlov in the beginning of the 20th century as the “what is that?” reflex, are elicited by novel innocuous stimuli and can be considered a the unit of attentional processing (Barry, 2009). The OR is a whole-body reflex, involving adjustments in a large range of physiological systems, including visuomotor reflexes, electrodermal activity, respiratory and cardiovascular changes, and changes in whole-brain activity (Sokolov, 1960, 1963a, 1963b). While innocuous stimuli elicit ORs, painful and threatening stimuli elicit a separate class of responses, the defense reflex (Sokolov, 1960, 1963a, 1963b). In spite of that, situations of potential or distal threat can lead to hypervigilance, in which innocuous stimuli can lead to defense reflexes (DRs). For example, in zebrafish (*Danio rerio*), exposure to an alarm substance – an olfactory stimulus that represents a distal threat – leads to a shift from ORs to DRs elicited by an innocuous visual stimulus (do Nascimento et al., 2020).

Arousal plays an important role in this OR-to-DR shift. The arousal level of an organism can serve as an “amplifying factor”, decreasing habituation (Sokolov, 1963a), or, at higher levels, lead to “false alarms” in a state of hypervigilance. The neural bases of this OR-to-DR shift probably involve the overlap between circuits for visual attention and defensive behavior. For example, in a preparation of eye-brain-spinal cord of lampreys (both *Lampetra fluviatilis* and *Petromyzon marinus*), recordings of the ventral roots were able to discriminate between a unilateral burst response corresponding to orientation of the head toward slowly expanding or moving stimuli, particularly within the anterior visual field, that corresponds to a OR, and a unilateral or bilateral burst response triggering avoidance in response to rapidly expanding looming stimuli or moving bars, corresponding to a DR (Suzuki et al., 2019). These fictive responses were enhanced by pharmacological blockade of brainstem-projecting neurons in the deep layers of the optic tectum (OT)(Suzuki et al., 2019). Similarly, in goldfish (*Carassius auratus*), unilateral tectal ablation abolishes the OR towards the contralateral hemifield, while electrical stimulation of the OT evokes contraversive eye movements (Torres et al., 2005). Habituation of ORs appear to depend on telencephalic input in goldfish, as electrical stimulation of the dorsomedial (Dm) and dorsocentral (Dc) regions of the telencephalon – the first being homologous to the associative amygdaloid nuclei of mammals (Maximino et al., 2013) – evoke cardiac and ventilatory responses, as well as bodily movements consistent with ORs (Quick & Laming, 1988). Likewise, telencephalic ablation decreased OR habituation in goldfish (Rooney & Laming, 1988), and animals with lesions in supracommissural (Vs) and postcommissural (Vp) regions of the telencephalon showed DRs to innocuous stimuli instead of ORs (Rooney & Laming, 1986). Thus, in fish, regions of the telencephalon that are part of the aversive brain system (do Carmo Silva, Lima-Maximino, et al., 2018) project to regions of the OT that organize ORs and DRs, modulating whether the animal will emit orienting or defense responses.

Both the deeper layers of the OT that project to the brainstem and the telencephalic regions involved in ORs and DRs show extensive serotonergic projections in fish (Lillesaar et al., 2009; Yokogawa et al., 2012) and mammals (Mooney et al., 1996). In rats, electrical stimulation of the superior colliculus (mammalian homologue of the OT) elicits head movements at lower current intensities and running and jumping responses at higher intensities; concurrent administration of the 5-HT reuptake blocker fluoxetine or the 5-HT1A receptor antagonist WAY100,635 decreases these responses, and administration of 5-HT in the midbrain, but not in the frontal cortex, produced the same effect (Dringenberg et al., 2003). In zebrafish larvae, a brief exposure to water flow elevates activity and sensitivity to behaviorally relevant visual cues that resemble an OR (Yokogawa et al., 2012). These effects are accompanied by increased activity in superior raphe (SR) neurons; genetic ablation of these cells abolished the increased sensitivity to visual cues (Yokogawa et al., 2012). In a similar paradigm, exposure to a conspecific alarm substance (CAS) induces a persistence hypervigilance in adult zebrafish, as indexed by lowered threshold to respond to stimuli of different sensory modalities; this state appears to be mediated by a synchronized state of neuronal dynamics in the dorsocentral and dorsodorsal telencephali (putative homologues of the mammalian cortex; Ganz et al., 2015; T. Mueller et al., 2011; Yáñez et al., 2022) and CAS leads to an intense activation of neurons in the SR (Zhao et al., 2024). Transgenic zebrafish lacking *tph2*, the rate-limiting enzyme in serotonin synthesis, do not show CAS-elicited neuronal synchrony nor hypervigilance after CAS (Zhao et al., 2024). Thus, serotonergic neurons of the SR appear to be involved in arousal-elicited amplifications in vigilance in zebrafish.

We have previously shown that an innocuous visual stimulus elicits an orienting response zebrafish, characterized by approaching the stimulus and projecting the body towards it (do Nascimento et al., 2020). We have also determined that, during the OR, the stimulus is investigated primarily with the left eye (do Nascimento et al., 2020), consistent with lateralization of novelty processing in this species (Miklósi et al., 1997). However, when animals were exposed to CAS – a mixture released in the water when club cells in the skin are damaged, signaling threat to conspecifics in a shoal (Maximino et al., 2019) –, this OR changes to a defense reflex, characterized by increased erratic swimming and escaping the stimulus. We have previously demonstrated a role for serotonin in other components of the alarm response: after exposure to CAS, extracellular 5-HT levels are elevated in zebrafish brains (Maximino et al., 2014) and monoamine oxidase activity is decreased (Lima-Maximino et al., 2020). Zhao et al. (2024) observed serotonin release in the dorsocentral telencephalon (Dc) after CAS exposure. Moreover, some components of the alarm response could be blocked by treatment with *para*-chlorophenylalanine (pCPA), a false substrate of tryptophan hydroxylase (TPH) which blocks serotonin synthesis, or metergoline, a non-selective antagonist at 5-HT receptors (Lima-Maximino et al., 2020). Similar effects were observed in transgenic zebrafish which do not express *tph2* (Zhao et al., 2024). The present work attempts to unravel the roles of 5-HT in this arousal-elicited OR-to-DR shift in adult zebrafish. In order to do that, animals were treated with either *para*-chlorophenylalanine (pCPA), a false substrate of tryptophan hydroxylase which blocks serotonin synthesis, or 5-hydroxytryptophan (5-HTP), a 5-HT precursor. It was hypothesized that pCPA-treated animals would display an attenuated effect of CAS, while 5-HTP would potentiate this effect.

## Methods

### Animals and housing

48 adult wild-type zebrafish from the longfin phenotype were used in the pCPA experiments, and 40 animals were used in the 5-HTP experiments. Sample sizes were not calculated prior to the experiments; however, *a posteriori* power calculations were made, and are reported with each statistical analysis. The populations used are expected to better represent the natural populations in the wild, due to its heterogeneous genetic background (Parra et al., 2009; Speedie & Gerlai, 2008). Animals were bought from a commercial vendor (Belém/PA) and collectively maintained in 40 L tanks for at least two weeks before the onset of experiments. The sex proportion was estimated to be of 1 male:1 female, as judged by body morphology; however, sex was not determined either through body nor gonadal morphology for each individual in the experiments, and therefore was not included as a variable. Animals were acquired from commercial vendors (Fernando Peixes, Belém/PA, and PisciculturaPower Fish, Itaguaí/RJ), and delivered to the laboratory with an approximate age of 3 months (standard length = 12.9 ± 1.6 mm), being quarantined for at least two weeks before experiments begun. Throughout experiments, animals were estimated to be 4 months old (standard length = 21.7 ± 3.1 mm) The animals were fed daily with fish flakes. The tanks were kept at constant temperature (28 °C), oxygenation, light cycle (14:10 LD photoperiod) and a pH of 7.0-8.0, according to standards of care for zebrafish (Lawrence, 2007). Animals were used for only one experiment to reduce interference from apparatus exposure. Potential suffering of animals was minimized by controlling for the aforementioned environmental variables. Furthermore, in the all experiments the animals used were handled, anesthetized and sacrificed according to the norms of the Brazilian Guideline for the Care and Use of Animals for Scientific and Didactic Purposes (CONCEA, 2017). Since, in all cases, animals were individually stressed and/or individually treated, experimental unit was a single animal, and therefore all sample sizes refer to number of animals. The experimental protocols were approved by the Animal Use Ethics Committee of UNIFESSPA (Universidade Federal do Sul e do Sudeste do Pará) protocol number 23479.019576/2022-43.

### Drug treatments

For drug treatment, animals were injected intraperitoneally with either vehicle (DMSO 10%), pCPA (150 mg/kg), or 5-HTP (5 mg/kg), in a repeated dosing schedule; animals were injected once per day for two days, followed by 24 h without drug injection, after which they were subjected to the behavioural tests. This treatment schedule is typically used in studies using pCPA to induce serotonin depletion in the central nervous system, and has been shown to reduce 5-HT levels in the zebrafish brain to ∼10% of controls (Maximino et al. 2013). After injection, animals were returned to collective tanks (maximum of 12 animals/tank); as a result, at least 4 different tanks were used per experiment, which helped to prevent pseudo-replication. Tank number was added in statistical analysis as a dummy variable, but showed no effect at p < 0.05. Animals were randomly allocated to groups using a random number generator (http://www.jerrydallal.com/random/random_block_size_r.htm), with each subject randomized to a single (vehicle/drug) treatment using random permuted blocks. For each experiment, animals were treated and tested in the order of allocation (i.e., randomly). In all experiments, experimenters and data analysts were blinded to drugs by using coded vials (with the same code used for randomization); blinding was removed only after data analysis.

### Exposure to alarm substance (CAS)

For the exposure stage, each animal was transferred individually to a container (2 L) where, after 3 minutes of acclimatization, it was carefully exposed to 7 ml of alarm substance (CAS), extracted using a standardized protocol (do Carmo Silva, Rocha, et al., 2018). Briefly, donor animals (size-matched to animals which were exposed to CAS) were cold-anesthetized, sacrificed by spinal section, and CAS was extracted by making 15 shallow cuts on each size of the carcass, avoiding drawing blood, and washing the cuts with 10 ml double distilled water. CAS from a single donor was used per animal, in a nominal concentration of 7 ml CAS/5 L water; CAS was always fresh-prepared, and delivered directly into the apparatus water. As a negative control, a group with the same amount of animals was exposed to the same volume of distilled water, according to the protocol of Lima-Maximino et al. (2020). The animals remained exposed for the entire session. All stages of the experiment were performed under constant white Gaussian noise, producing an average of 58 dB above the tank. Light levels above the tanks were measured using a handheld light meter, and ranged from 251 to 280 lumens (coefficient of variation = 3.399% between subjects).

Animals were randomly allocated to groups using a random number generator (http://www.jerrydallal.com/random/random_block_size_r.htm), with each subject randomized to a single treatment using random permuted blocks. For each experiment, animals were treated and tested in the order of allocation (i.e., randomly). In all experiments, experimenters and data analysts were blinded to drugs and treatment by using coded vials (with the same code used for randomization); thus, results were coded for both drug treatment (using randomly assigned letters for each drug) and exposure (using randomly assigned numbers for either water or CAS). Blinding was removed only after data analysis. Experiments were always run between 08:00AM and 02:00 PM. After experiments, animals were sacrificed by prolonged bath in ice-cold water (< 12 °C), followed by spinal transection (Matthews & Varga, 2012).

### Stimulus presentation

A blue circle (R: 0; G: 0; B: 128; maximum 256) was used, based on the ability of zebrafish to discriminate blue from green stimuli in a simple discrimination paradigm (K. P. Mueller & Neuhauss, 2012). Stimulus size was 72° of horizontal visual angle, calculated considering the diameter of the stimulus in the screen when it is at the front of the animal, and the animal is in the centre of the tank. The size was based on previous work which show that this size is insufficient to produce an escape response in adult zebrafish that were not exposed to CAS (do Nascimento et al., 2020).

Animals were transferred to the tank and left to acclimate for 3 min before the presentation of the stimuli. The computer screen was turned on for the entire experiment, including acclimatization time. Using an animation based on LibreOffice Impress (v. 6.0.7.3), the stimulus was presented in 1-min intervals (“trials”) interspersed with 1-min stimulus-free periods. In trials in which the stimulus was on, the stimulus was presented for the entire duration of the trial. 10 trials were made; total session length was 22 minutes per animal. The number of trials and intertrial durations were chosen based on previous work on arousal in goldfish (Laming & Savage, 1980).

### Quantification of orienting and defensive responses

The experimental setup consisted of two 13 × 13 × 17 (depth × length × depth cm each) glass tanks, positioned side by side, and filled with 2 L tank water. A mobile phone (Redmi Note® 10S, camera resolution 1080p at 30 fps) was positioned above the tanks to allow recording and offline behavioral scoring. All focal fish behaviors were therefore tracked from a top-down perspective, using both TheRealFishTracker (http://www.dgp.toronto.edu/~mccrae/projects/FishTracker/) and fishtracker (https://github.com/joseaccruz/fishtracker). For each behavioral video, a 2D region (arena) was defined for tracking, comprising the inner area of the tank. A region-of-interest (ROI) of 13 x 8.5 cm was determined as “stimulus zone”. The following variables were recorded, using TheRealFishTracker:

- Total time in the stimulus zone (s);
- Absolute turn angle (°), a measure of erratic swimming (Lima-Maximino et al., 2020);
- Speed (cm/s)

Following Abril-de-Abreu, Cruz, & Oliveira (2015), a region of interest (ROI) of 13 x 3 cm (approximately 25% of the tank), corresponding to the width of the tank and the mean body length of an adult zebrafish, was defined in the area of the arena closer to the computer screen in fishtracker. Animal orientation is defined as a mean projection vector *R_proj_* (Abril-de-Abreu et al., 2015), determined by the transformation of each orientation angle in a vector *r_i_*= (*cosα_i_, sinα_i_*), where αi represents the angle formed by the direction of the centroid-head axis relative to the horizontal axis in each frame. The resulting mean vector, with length R = ||r||, is a measure of directional focus, and varies between 0 and 1. The projection of this vector R in the stimulus direction (arbitrarily defined as 180°) is defined as *R_proj_* = *-Rcosα*. Since *R_proj_*is a mean vector – that is, it takes projection angles in each frame and averages then through the whole 1-min trial –, if the animal is consistently orienting throughout most of the trial, values tend to be positive; if the animal is consistently facing away from the stimulus, values tend to be negative; and if the animal is orienting towards many different directions during the trial, values tend to be close to 0. Thus, positive values indicate direction towards the stimulus, while negative values indicate direction away from the stimulus, and values close to 0 indicate absence of orienting.

Eye use was estimated by analysing projection angles averaged across trials in which the stimulus was on and trials in which the stimulus was off, with angles between 150° and 179° recorded as “left eye use” (“L”), angles between 181° and 210° recorded as “right eye use” (“R”), and average angles above 210° and below 150° recorded as “no preference”. Thus, eye use was defined as orienting the head in an angle of up to 30°, consistent with what is observed in other experiments on eye use in zebrafish (do Nascimento et al., 2020; Miklósi et al., 1997; Miklósi & Andrew, 1999).

### Statistical analysis

For all data except directional angle (*R*), three-way (treatment X drug X trial type) analyses of variance were applied. When p-values were smaller than 0.05, Tukey’s HSD post-hoc tests were applied. For directional angle data, circular means were analyzed using projected normal circular generalized linear models; data were compared based on the posterior distribution of the circular mean for each condition, based on lower and upper intervals of the highest posterior density interval (HPD)(Cremers & Klugkist, 2018). Data and scripts can be found at https://github.com/lanec-unifesspa/5-HT-CAS/tree/master/orienting.

## Results

### Experiment 1

Significant main effects of trial type, treatment, and drug were found for the time near the stimulus (Table 1; Figure 1A). In general, animals spent almost half of the each 1-min. trial near the stimulus when it was turned on (24.07 ± 15.78 s with stimulus on vs. 12.54 ± 8.65 s with stimulus off [mean ± S.D.]); thus, when the stimulus was turned on, animals approached it. Significant interactions were also found between drug and treatment, drug and trial type, treatment and trial type, and a three-way interaction between all variables (Table 1). Post-hoc Tukey’s HSD tests found that the approach response to the stimulus was abolished in the presence of CAS (d = 1.13, p < 0.001); in fact, it is apparent that the approach response is turned into an escape response in the presence of CAS, as individuals in this condition spend less time near the stimulus when it was turned on than when it was turned off. pCPA abolished this effect, as individuals treated with the drug and exposed to CAS spent more time near the stimulus when it was turned on than when it was turned off (d = -4.81, p < 0.001). In summary, when the stimulus was turned on, control animals approached it, and CAS-exposed animals escaped it, and this last effect was prevented by pCPA.

**Figure 1.**
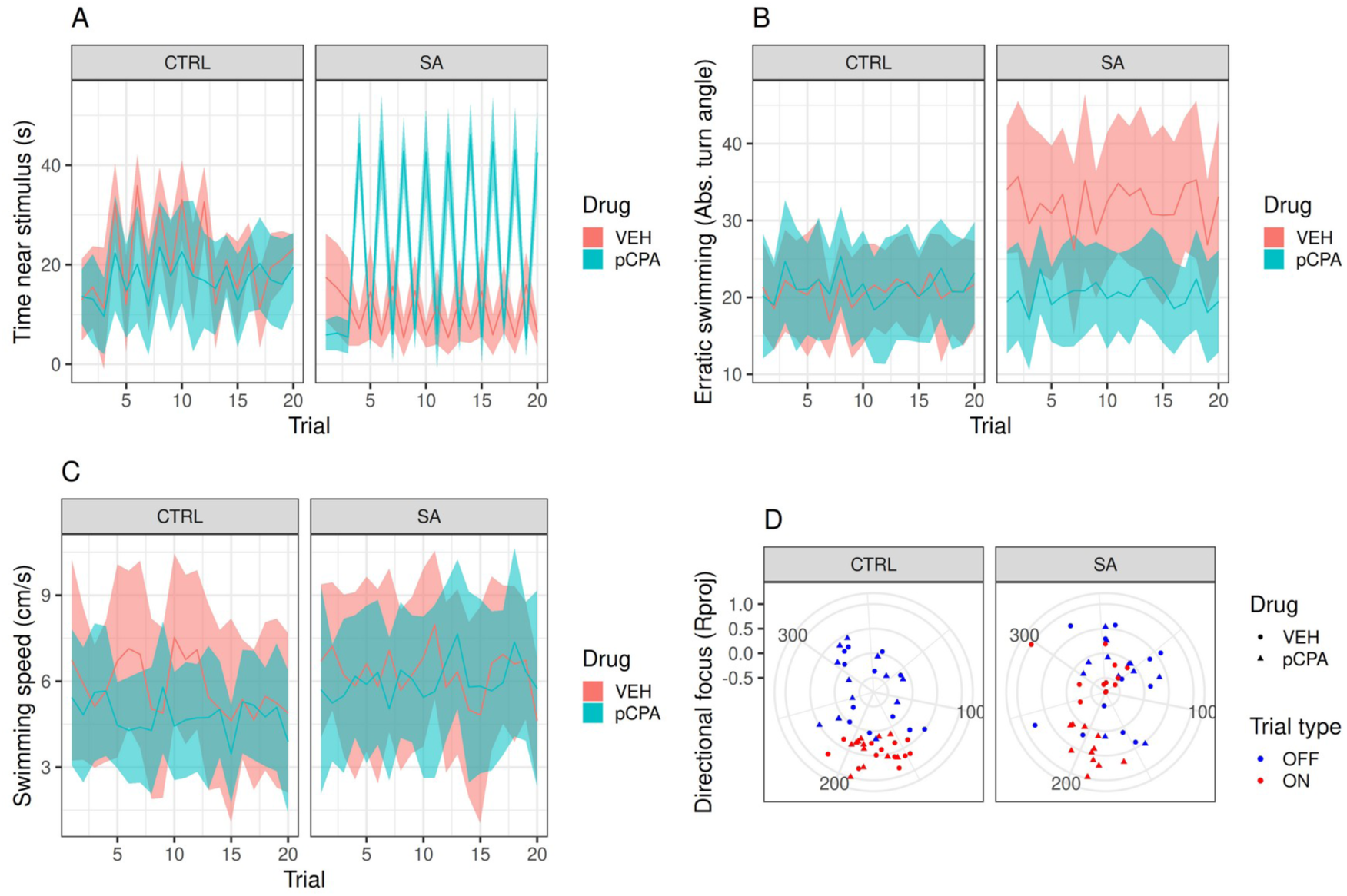
pCPA blocked CAS-elicited OR-to-DR shift. (A) Presenting zebrafish with a visual stimulus elicits approach to the stimulus zone, as shown by peaks in the time near stimulus in trials in which the stimulus was on. CAS inverts this approach response, decreasing time near the stimulus when the stimulus is turned on. pCPA treatment prevents the effects of CAS. (B) The visual stimulus does not alter erratic swimming, but CAS increases it, an effect that is attenuated by pCPA. (C) No effects of the visual stimulus, CAS, or pCPA were observed on swimming speed. (D) Zebrafish project their bodies towards the stimulus (position indicated by the letter “S”) when it is on, while CAS leads animals to project their body away from the stimulus; pCPA abolishes the effects of CAS on the OR. In (A-C), red lines represent group averages for vehicle-treated animals, with pink ribbons representing 95% confidence intervals; blue lines represent group averages for pCPA-treated animals, with light blue ribbons representing 95% confidence intervals. In (D), each dot represents average values for a single individual, with the distance from the center of the polar plot representing the directional focus *R_proj_*, and its position in the circular plot represents the angle (in °) of orientation; both variables are average across trial types (on vs. off).

**Table 1.** Summary of statistical analysis on the variables *time near stimulus*, *erratic swimming*, and *directional focus* for Experiment 1.

| Endpoint | Independent variable | df | F | p | $\omega^2$ | Power |
| --- | --- | --- | --- | --- | --- | --- |
| Time near stimulus | Trial type (on vs. off) | 1, 949 | 514.88 | < 0.001 | 0.17 | 0.98 |
|  | Treatment (CTRL vs. CAS) | 1, 949 | 11.3 | < 0.001 | 0.00 | 0.1 |
|  | Drug (vehicle vs. pCPA) | 1, 949 | 148.57 | < 0.001 | 0.05 | 0.78 |
|  | Treatment x Drug | 1, 949 | 303.64 | < 0.001 | 0.10 | 0.97 |
|  | Trial type x Drug | 1, 949 | 369.29 | < 0.001 | 0.13 | 0.99 |
|  | Trial type x Treatment | 1, 949 | 32.11 | < 0.001 | 0.01 | 0.22 |
|  | Trial type x Treatment x Drug | 1, 949 | 664.12 | < 0.001 | 0.22 | 0.99 |
| Erratic swimming | Trial type | 1, 949 | 0.81 | 0.369 | -0.01 | 0.1 |
|  | Treatment | 1, 949 | 133.83 | < 0.001 | 0.10 | 0.53 |
|  | Drug | 1, 949 | 95.9 | < 0.001 | 0.07 | 0.11 |
|  | Treatment x Drug | 1, 949 | 152.03 | < 0.001 | 0.11 | 0.74 |
|  | Trial type x Drug | 1, 949 | 1.94 | 0.164 | 0.0 | 0.05 |
|  | Trial type x Treatment | 1, 949 | 0.07 | 0.785 | -0.0 | 0.05 |
|  | Trial type x Treatment x Drug | 1, 949 | 0.03 | 0.868 | -0.0 | 0.05 |
| Swimming speed | Trial type | 1, 949 | 0.01 | 0.918 | -0.0 | 0.05 |
|  | Treatment | 1, 949 | 8.11 | 0.004 | 0.01 | 0.05 |
|  | Drug | 1, 949 | 6.63 | 0.01 | 0.01 | 0.05 |
|  | Treatment x Drug | 1, 949 | 2.39 | 0.123 | 0.0 | 0.05 |
|  | Trial type x Drug | 1, 949 | 0.7 | 0.403 | -0.0 | 0.05 |
|  | Trial type x Treatment | 1, 949 | 0.99 | 0.321 | -0.0 | 0.05 |
|  | Trial type x Treatment x Drug | 4, 949 | 0.54 | 0.461 | -0.0 | 0.05 |
| Directional focus ($R_{proj}$) | Trial type | 1, 88 | 5.89 | 0.017 | 0.03 | 0.43 |
|  | Treatment | 1, 88 | 6.76 | 0.011 | 0.03 | 0.26 |
|  | Drug | 1, 88 | 3.42 | 0.068 | 0.01 | 0.15 |
|  | Treatment x Drug | 1, 88 | 8.91 | 0.004 | 0.05 | 0.37 |
|  | Trial type x Drug | 1, 88 | 22.54 | < 0.001 | 0.11 | 0.53 |
|  | Trial type x Treatment | 1, 88 | 24.87 | < 0.001 | 0.14 | 0.84 |
|  | Trial type x Treatment x Drug | 4, 88 | 20.81 | < 0.001 | 0.11 | 0.74 |

Significant main effects of treatment, and drug were found for the erratic swimming (Table 1; Figure 1B). In general, animals exposed to CAS showed higher erratic swimming than non-exposed individuals at both trial types (20.87° ± 6.66 for controls vs. 26.49° ± 9.51 for CAS-exposed animals), producing a medium-sized effect (d = - 0.75, p < 0.001). A significant interaction between drug and treatment was also found (Table 1). Vehicle-treated, CAS-exposed animals showed increased erratic swimming in relation to controls (d = -1.56, p < 0.001), while pCPA-treated animals did not show differences in relation to controls when not exposed to CAS (d = -0.12, p = 0.59), and showed lower erratic swimming in relation to vehicle-treated, CAS-exposed animals (d = 1.44, p < 0.001). In summary, CAS increased erratic swimming, an effect that was prevented by pCPA and independent of stimulus presentation.

Significant effects of treatment and drug were found for swimming speed, but interaction effects were absent (Table 1, Figure 1C). Treatment with pCPA generally decreased swimming speed (d = 0.17, p = 0.01), while exposure to CAS increased it (d = -2.85, p = 0.004).

By examining the lower and upper bounds of the 95% highest posterior density interval (HPD) in Table 2, it is apparent that the HPD intervals of the circular mean of all conditions overlap, and thus we conclude that there is not enough evidence to reject the null hypothesis that the circular means do not differ and that there is no effect of treatment, drug, or trial type on the average directional angle (Figure 1D). In summary, stimulus presentation (on trials) elicited an orientation towards the stimulus in control animals and away from it in CAS-exposed animals, and this latter effect was prevented by pCPA.

**Table 2.** Posterior estimates of the circular means of the projection angle for Experiment 1.

| Treatment | Drug | Trial type | Mode | Mean ± S.D. | LB HPD | UB HPD |
| --- | --- | --- | --- | --- | --- | --- |
| CTRL | VEH | off | 1.873 | 1.858 ± 0.849 | -0.214 | 3.779 |
| | | on | 0.321 | 0.492 ± 2.282 | -2.278 | 3.574 |
| CAS | VEH | off | 2.156 | 1.47 ± 2.731 | -2.322 | 3.652 |
| | | on | -0.157 | 1.203 ± 2.503 | 2.714 | 2.284 |
| CTRL | pCPA | off | 2.662 | 2.797 ± 2.66 | -1.146 | 4.701 |
| | | on | 1.774 | 1.256 ± 2.662 | -1.521 | 4.343 |
| CAS | pCPA | off | 3.201 | 2.115 ± 2.547 | 0.137 | 6.035 |
| | | on | 1.021 | 1.246 ± 2.513 | -1.689 | 4.126 |

Significant main effects of trial type and treatment were found for directional focus (Table 1, Figure 1D). Significant treatment X drug, trial type X drug, trial type X treatment, and three-way interactions were also found (Table 1). In general, directional focus increased when the stimulus was turned on (d = -0.5, p = 0.017). When vehicle-injected animals were exposed to CAS, directional focus was negative when the stimulus was turned on (d = 1.7, p = 0.002 vs. stimulus off; d = 3.09, p < 0.00 vs. CTRL). Treatment with pCPA abolished this effect.

Finally, significant associations between eye use and trial type, treatment, and drug were found (χ²[df = 14] = 85.033, p < 0.0001). No association was found when the stimulus was turned off, but significant associations were found when the stimulus was turned on. In these latter cases, control-exposed individuals treated with vehicle used more the left eye, and control-exposed individuals treated with pCPA used more the right eye; CAS-exposed individuals treated with vehicle showed no eye preference, and CAS-exposed individuals treated with pCPA used more the left eye. In summary, control individuals preferred to inspect the stimulus with the left eye, while pCPA switches this preference for the right eye. pCPA also prevented the tendency in CAS-exposed individuals to not display eye preferences.

### Experiment 2

Main effects of trial type, treatment, and drug were found for the time spent near the stimulus zone in Experiment 2 (Table 3, Fig 1A). Interactions between stimulus type and treatment, drug and treatment, and three-way interactions were found. In general, CAS-exposed animals treated with vehicle showed decreased time in the stimulus area when the stimulus was turned on (d = 0.6, p = 0.003), while control animals treated with vehicle showed more time in the stimulus area when the stimulus was turned on (d = - 0.93, p < 0.001). Again, presentation of the stimulus elicited an approach response in control animals, and an escape response in CAS-exposed animals. This effect was attenuated by treatment with 5-HTP (d =-0.4, p = 0.356 vs. CAS-exposed animals with stimulus on); however, 5-HTP also appeared to abolish the OR in animals which were not exposed to CAS (d = 0.18, p = 0.93, CTRL animals treated with 5-HTP, stimulus off vs. on). In summary, while the effects of trial type and CAS from Experiment 1 were replicated, the effects of 5-HTP on both controls and CAS-exposed individuals is consistent with an attenuation in responsiveness.

**Table 3.** Summary of statistical analysis on the variables *time near stimulus*, *erratic swimming*, and *directional focus* for Experiment 2.

| Endpoint | Independent variable | df | F | p | $\omega^2$ | Power |
| --- | --- | --- | --- | --- | --- | --- |
| Time near stimulus | Trial type | 1, 664 | 16.28 | < 0.001 | 0.02 | 0.13 |
|  | Treatment | 1, 664 | 48.71 | < 0.001 | 0.07 | 0.36 |
|  | Drug | 1, 664 | 23.81 | < 0.001 | 0.03 | 0.22 |
|  | Treatment x Drug | 1, 664 | 4.95 | 0.026 | 0.01 | 0.05 |
|  | Trial type x Drug | 1, 664 | 3.11 | 0.078 | 0.0 | 0.05 |
|  | Trial type x Treatment | 1, 664 | 181.65 | < 0.001 | 0.21 | 0.85 |
|  | Trial type x Treatment x Drug | 1, 664 | 8.76 | 0.003 | 0.01 | 0.05 |
| Erratic swimming | Trial type | 1, 606 | 0.19 | 0.662 | -0.0 | 0.05 |
|  | Treatment | 1, 606 | 17.2 | < 0.001 | 0.03 | 0.18 |
|  | Drug | 1, 606 | 2.93 | 0.087 | 0.0 | 0.05 |
|  | Treatment x Drug | 1, 606 | 8.15 | 0.004 | 0.01 | 0.05 |
|  | Trial type x Drug | 1, 606 | 0.09 | 0.767 | -0.0 | 0.05 |
|  | Trial type x Treatment | 1, 606 | 0.07 | 0.785 | -0.0 | 0.05 |
|  | Trial type x Treatment x Drug | 1, 606 | 0.01 | 0.934 | -0.0 | 0.05 |
| Swimming speed | Trial type | 1, 685 | 2.21 | 0.138 | 0.0 | 0.05 |
|  | Treatment | 1, 685 | 15.38 | < 0.001 | 0.02 | 0.13 |
|  | Drug | 1, 685 | 78.9 | < 0.001 | 0.09 | 0.49 |
|  | Treatment x Drug | 1, 685 | 70.55 | < 0.001 | 0.08 | 0.45 |
|  | Trial type x Drug | 1, 685 | 1.0 | 0.317 | 0.0 | 0.05 |
|  | Trial type x Treatment | 1, 685 | 1.52 | 0.217 | 0.0 | 0.05 |
|  | Trial type x Treatment x Drug | 1, 685 | 0.0 | 0.97 | -0.0 | 0.05 |
| Directional focus ($R_{proj}$) | Trial type | 1, 88 | 5.502 | 0.0212 | 0.06 | 0.58 |
|  | Treatment | 1, 88 | 3.033 | 0.085 | 0.03 | 0.31 |
|  | Drug | 1, 88 | 25.501 | < 0.001 | 0.225 | 0.98 |
|  | Treatment x Drug | 1, 88 | 1.049 | 0.309 | 0.012 | 0.15 |
|  | Trial type x Drug | 1, 88 | 14.964 | < 0.001 | 0.145 | 0.96 |
|  | Trial type x Treatment | 1, 88 | 6.091 | 0.015 | 0.065 | 0.62 |
|  | Trial type x Treatment x Drug | 1, 88 | 0.031 | 0.861 | 0.0 | 0.05 |

Main effects of treatment, but not drug nor trial type, were found for erratic swimming (Table 3, Figure 2B). Drug X treatment interactions were also found. In general, CAS-exposed animals had lower erratic swimming than non-exposed animals (d = 0.34, p < 0.001), an effect that appears to be mainly driven by the higher levels of erratic swimming in non-exposed, 5-HTP-treated animals regardless of stimulus type.

**Figure 2.**
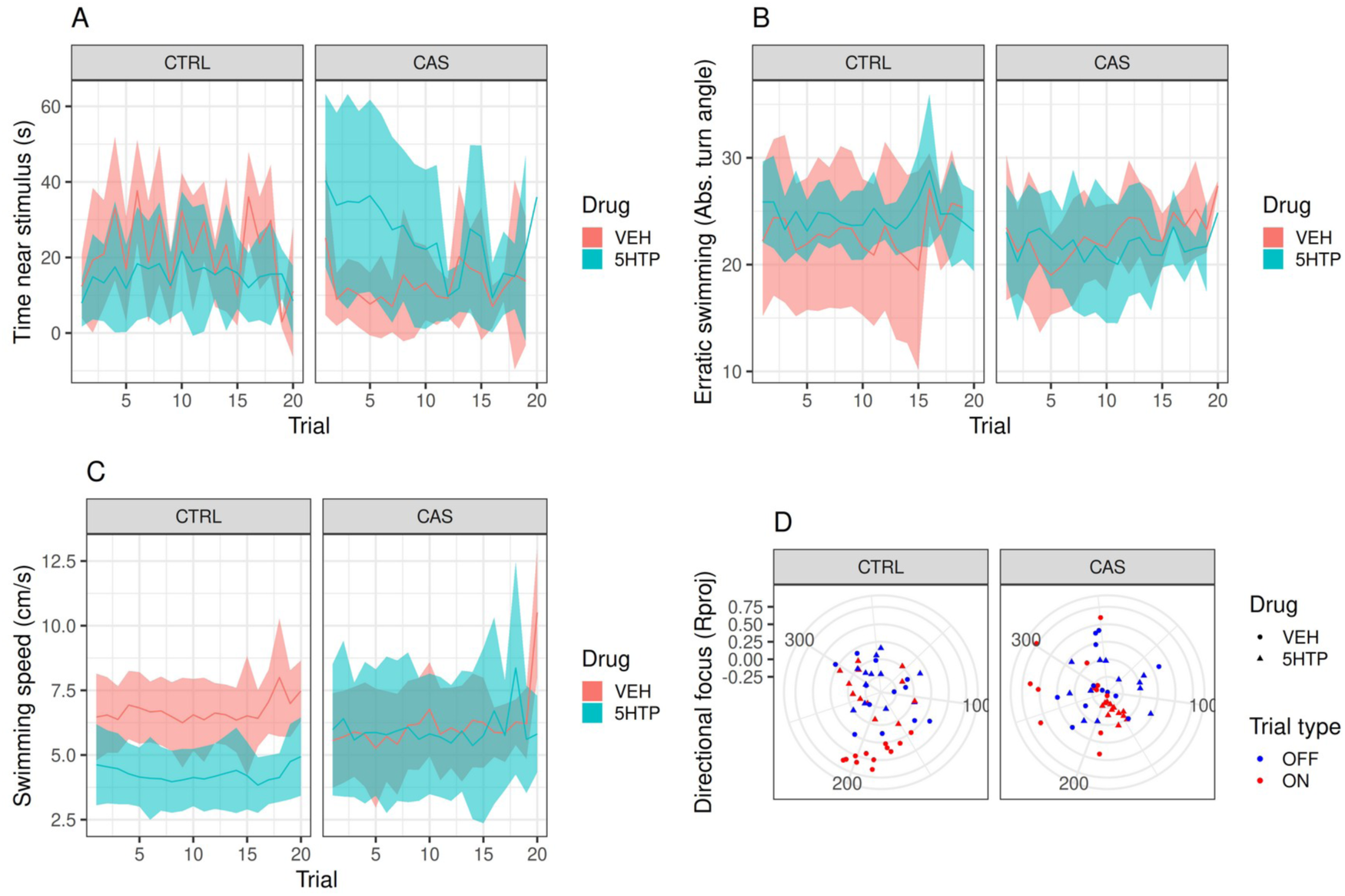
5-HTP attenuates OR responses in control zebrafish. (A) Presenting zebrafish with a visual stimulus elicits approach to the stimulus zone, as shown by peaks in the time near stimulus intrials in which the stimulus was on. CAS inverts this approach response, decreasing time near the stimulus when the stimulus is turned on. 5-HTP treatment decreases the approach in both control and CAS-exposed zebrafish. (B) The visual stimulus does not alter erratic swimming, but CAS increases it, an effect that is not altered by 5-HTP. (C) No effects of the visual stimulus or CAS were observed on swimming speed, but 5-HTP treatment decreased swimming speed. (D) Zebrafish project their bodies towards the stimulus (position indicated by the letter “S”) when it is on, while CAS leads animals to project their body away from the stimulus; 5-HTP attenuates or abolishes this OR. In (A-C), red lines represent group averages for vehicle-treated animals, with pink ribbons representing 95% confidence intervals; blue lines represent group averages for pCPA-treated animals, with light blue ribbons representing 95% confidence intervals. In (D), each dot represents average values for a single individual, with the distance from the center of the polar plot representing the directional focus *R_proj_*, and its position in the circular plot represents the angle (in °) of orientation; both variables are average across trial types (on vs. Off).

Significant main effects of drug and treatment were found for swimming speed (Table 3, Figure 2C). Significant interactions were also found between drug and treatment. 5-HTP-treated animals showed consistently lower swimming speeds than all other groups, regardless of stimulus type.

Main effects of trial type and drug were found for directional focus (Table 4), and interactions between trial type and drug and trial type and treatment were also found. By examining the lower and upper bounds of the 95% highest posterior density interval (HPD) in Table 4, no overlap in the HPD intervals of the circular mean of on and off trials in CTRL animals (both treated with vehicle and 5-HTP), and thus we conclude that, for animals not exposed to CAS, the average directional angles were different in these conditions, with the body directed towards the stimulus in on trials (Figure 2D).

**Table 4.** Posterior estimates of the circular means of the directional angle for Experiment 2.

| Treatment | Drug | Trial type | Mode | Mean ± S.D. | LB HPD | UB HPD |
| --- | --- | --- | --- | --- | --- | --- |
| CTRL | VEH | off | -1.543 | -1.989 ± 2.574 | 0.522 | 0.066 |
| | | on | -1.449 | -1.305 ± 0.616 | -2.403 | 0.004 |
| CAS | VEH | off | -2.147 | -2.126 ± 0.24 | -2.6 | -1.62 |
| | | on | -0.703 | -0.782 ± 0.745 | -2.321 | 0.698 |
| CTRL | 5-HTP | off | -2.414 | -2.552 ± 1.34 | 0.257 | 5.70 |
| | | on | -0.638 | -0.677 ± 0.653 | -2.009 | 0.68 |
| CAS | 5-HTP | off | -1.61 | -1.57 ± 1.49 | 2.11 | 1.35 |
| | | on | -2.349 | -2.795 ± 1.265 | 0.891 | 5.987 |

Finally, significant associations between eye use and trial type, treatment, and drug were found (χ²[df = 14] = 23.87, p < 0.0001). No association was found when the stimulus was turned off, but significant associations were found when the stimulus was turned on. In these cases, control-exposed individuals treated with vehicle used more the left eye, and control-exposed individuals treated with 5-HTP lost this preference; CAS-exposed individuals treated with vehicle showed no eye preference, and CAS-exposed individuals treated with 5-HTP used more the right eye.

## Discussion

The present work tested the participation of serotonin on orienting responses of zebrafish, as well as in the shift from orienting response to defense reaction (OR-to-DR shift) elicited by a conspecific alarm substance (CAS). pCPA blocked CAS-elicited OR- to-DR shift, as shown both by the shift from an approach to an escape response when the stimulus was turned on and by the changes in directional focus. pCPA had no effects on erratic swimming; importantly, erratic swimming was not significantly altered by trial type, only by CAS, suggesting again that this stimulus induces an aversive state that is unrelated to visual stimulation, while simultaneously leading to the OR-to-DR shift. 5-HTP, on the other hand, attenuated the CAS-elicited OR-to-DR shift and abolished ORs in non-exposed animals, suggesting that the serotonergic tone is more important in regulating the general arousal levels. These results are consistent with the prediction that pCPA would attenuate the OR-to-DR shift, but not the prediction that 5-HTP would potentiate it.

The OR observed in the present experiments essentially replicates previous findings from our group (do Nascimento et al., 2020). The OR was characterized by an approach response (more time spent near the stimulus when it was turned on) and a body oriented towards the stimulus (higher directional focus), with predominant use of the left eye to inspect the stimulus. We also replicated our previous observation that CAS elicits an OR-to-DR shift in zebrafish (do Nascimento et al., 2020). This shift was characterized by avoiding the stimulus zone when the visual stimulus was turned off and reduced directional focus. These effects are consistent with a defensive reaction, with CAS leading to an attentional bias in which the animal “misinterprets” the innocuous visual stimulus as a threatening stimulus. Evidence for an aversive state elicited by CAS is also present at the increased, stimulus-independent elevation in erratic swimming.

Importantly, we found that pCPA abolished the OR-to-DR shift in CAS-exposed animals. This effect was also accompanied by the abolishing of the effect of CAS on erratic swimming, suggesting that, while the serotonergic tone is permissive to ORs elicited by visual non-threatening stimuli, it is also necessary to mount a defensive response. This is consistent with previous findings with zebrafish, in which pCPA blocked the effects of CAS on responses to distal and proximal threat (Lima-Maximino et al., 2020). These findings are also consistent with previous work which shows that transgenic zebrafish that do not express *tph2* are not sensitive to arousal-eliciting effects of CAS (Zhao et al., 2024). The OR-attenuating effect of pCPA was also observed in rats, in which the drug significantly reduced the responding to the first presentation of a tone, but did not alter the rate of habituation (File, 1975). On the other hand, pCPA has been shown to enhance startle responses in rats (Carlton & Advokat, 1973), suggesting that the role of serotonin in orienting and defensive responses can be opposite.

We initially hypothesized that 5-HTP would produce an opposite effect of pCPA in the OR-to-DR shift. However, we found that 5-HTP attenuated both the OR (in control animals) and the OR-to-DR shift (in CAS-exposed animals). 5-HTP also decreased swimming speed across treatments and exposure regimens, suggesting either a psychomotor effect or decreased responsiveness. Indeed, serotonin has been shown to suppress locomotion in zebrafish (Gabriel et al., 2009; Tran et al., 2016). An hypothesis to explain these effects is that the vigilance-promoting effects of serotonin are dependent on baseline levels of this transmitter, and increasing these levels above a certain threshold do not promote more arousal. In rhesus macaques, 5-HTP increases attention in animals that exhibit low baseline attention, while decreasing attention in animals that exhibit high baseline attention; importantly, this effect is associated with baseline concentrations of 5-HTP in the cerebrospinal fluid, with high baseline attention being associated with lower 5-HTP levels (Weinberg-Wolf et al., 2018). While our design does not allow to separate between baseline and post-drug effects, it is possible that the serotonergic tone is permissive to ORs elicited by non-threatening visual stimuli, while phasic stimulation of serotonin are less important in that regard. We have previously suggested that, for threatening stimuli, phasic serotonergic signals inhibit fear-like responses, while the serotonergic tone tracks aversive expectation values (Lima-Maximino et al., 2020). A similar mechanism might be involved in tracking novelty and expectation values in orienting responses.

Surprisingly, pCPA affected eye use during the presentation of the stimulus. Our results replicate previous results which show preference for the left eye in control animals during the presentation of the stimulus, while CAS-exposed animals showed no preference (do Nascimento et al., 2020). Moreover, we found that non-exposed individuals treated with pCPA showed a preference for the right eye when viewing the stimulus, while non-exposed animals treated with 5-HTP lost preference. When exposed to CAS, pCPA-treated animals shifted preference to the left eye, while 5-HTP-treated animals used more the right eye. In zebrafish, novel objects were investigated mainly with right frontal field, while in a second trial left frontal viewing tended to be used instead (Miklósi et al., 1997; Miklósi & Andrew, 1999). This is unlikely to be due to asymmetry in the habenular nuclei, which has been linked to preference in eye use (Dadda et al., 2010), because habenular projections to zebrafish raphe are not asymmetrical (Amo et al., 2010). Miklósi et al. (1997) suggested that the right eye is used when it is necessary to inhibit premature response in conditions of decision-making, and the left is used when it is necessary to track familiar objects. While we were not able to ascertain whether eye use changed during trials, it is possible that serotonin helps to track novelty, which could explain the effects of pCPA and 5-HTP on this variable.

Despite our findings that pCPA and 5-HTP altered the CAS-elicited OR-to-DR shift, a more thorough investigation of the specific mechanisms by which phasic and tonic serotonin modulate arousal and defensive responses to non-threatening visual stimuli is still needed. Given the high number of different serotonin receptors and the wide distribution of serotonergic innervation across the brain, the specific participation of receptors and brain regions is far from known. Nonetheless, these results point to an important serotonergic regulation of an emotion-cognition interface in vertebrates, which could represent a mechanism through which individuals respond to threats.

## Acknowledgments

This work was supported by a Conselho Nacional de Desenvolvimento Científico e Tecnológico (CNPq/Brazil) grant (Edital Universal 2016, #400726/2016-6).

